# Prediction of multiple conformational states by combining sequence clustering with AlphaFold2

**DOI:** 10.1101/2022.10.17.512570

**Authors:** Hannah K. Wayment-Steele, Sergey Ovchinnikov, Lucy Colwell, Dorothee Kern

## Abstract

AlphaFold2 (AF2) has revolutionized structural biology by accurately predicting single structures of proteins and protein-protein complexes. However, biological function is rooted in a protein’s ability to sample different conformational substates, and disease-causing point mutations are often due to population changes of these substates. This has sparked immense interest in expanding AF2’s capability to predict conformational substates. We demonstrate that clustering an input multiple sequence alignment (MSA) by sequence similarity enables AF2 to sample alternate states of known metamorphic proteins, including the circadian rhythm protein KaiB, the transcription factor RfaH, and the spindle checkpoint protein Mad2, and score these states with high confidence. Moreover, we use AF2 to identify a minimal set of two point mutations predicted to switch KaiB between its two states. Finally, we used our clustering method, AF-cluster, to screen for alternate states in protein families without known fold-switching, and identified a putative alternate state for the oxidoreductase DsbE. Similarly to KaiB, DsbE is predicted to switch between a thioredoxin-like fold and a novel fold. This prediction is the subject of future experimental testing. Further development of such bioinformatic methods in tandem with experiments will likely have profound impact on predicting protein energy landscapes, essential for shedding light into biological function.

## Introduction

Understanding the functions of any protein requires first understanding the complete set of conformational substates it can adopt^1^. For any protein structure prediction method, the task of predicting ensembles can be considered into two parts: an ideal method would (1) generate conformations encompassing the complete landscape and (2) score these conformations in accordance with the underlying Boltzmann distribution. The unprecedented accuracy of AlphaFold2 (AF2)^2^ at single-structure prediction has garnered interest in its ability to predict multiple conformations of proteins, yet AF2 has been demonstrated to fail in predicting multiple structures of metamorphic proteins^3^ and proteins with apo/holo conformational changes^4^ using its default settings.

Metamorphic proteins, or proteins that occupy more than one distinct secondary structure as part of their biological function^5–7^, are a useful set of model proteins to develop methods for predicting conformational ensembles, as they undergo particularly striking conformational changes. For instance, although the metamorphic protein KaiB only contains 108 residues, it undergoes a conformational shift that affects the secondary structure of roughly 40 residues in its C-terminal part, switching between a canonical thioredoxin-like structure and a unique alternate conformation ^8^. Fewer than 10 metamorphic protein families have been thoroughly experimentally characterized^5^, but those that have span a diverse range of functions. Fold-switching in proteins governs transcription regulation (RfaH in *E. coir*^9,10^), circadian rhythms (KaiB in cyanobacteria^8^), enzymatic activity (the selecase metallopeptidase in *M. janaschii*^11^), cell signaling (the chemokine lymphotactin in humans^12^), and cell cycle checkpoints (Mad2 in humans^13–16^). A computational analysis of the PDB that identified changes in secondary structure between protein models sharing the same sequence suggested that between 0.5-4% of all proteins are fold-switching^17^. The development of systematic methods to identify fold-switching proteins would aid in identifying new fold-switching proteins, highlight new structures and interactions to target for therapeutics^5^, as well as illuminate broader principles of protein structure, function, and evolutionary history that underlie known and as-of-yet undiscovered metamorphic proteins.

Despite demonstrations of AF2’s shortcomings in predicting several types of conformational changes in its default settings, Del Alamo et al. have demonstrated that subsampling the input MSA allows AF2 to predict conformational changes of an ion channel ^18^. Additionally, older coevolutionary-based methods ^19–22^ were able to extract multiple states of proteins with conformational changes, including ion channels ^19^, ligand-induced conformational changes ^23^, and multimerization-induced conformational changes ^24^. This line of research indicates that coevolutionary signals can be present for multiple conformational states, but computational methods must first deconvolve the signal from multiple states. Methods proposed to deconvolve signal when prior knowledge about one or more states is known include ablating signal from a known dominant state^25^ and supplementing the original multiple sequence alignment (MSA) with proteins known to occupy a rarer state^26^. However, simply subdividing a MSA and making predictions for portions of the MSA has also been used to detect variations in coevolutionary signal within a protein family^24,27,28^.

We hypothesized that metamorphic proteins might also demonstrate coevolutionary signal for multiple conformational states, and that if we could deconvolve this signal and input it separately into AF2, AF2 might be able to predict multiple conformations with high structural accuracy. We demonstrate that a simple MSA subsampling method – clustering sequences by sequence similarity – allows AF2 to predict both states of the metamorphic proteins KaiB, RfaH, and Mad2. Importantly, AF2 both samples the alternate structures and scores them with high confidence. Using KaiB as an example system, we further investigate the nature of AF2’s prediction of multiple states, as well as the biochemical underpinnings of its fold-switching. By comparing predictions from AF2 and the deep learning model MSA Transformer^29^, we posit that AF2’s prediction of one state comes from AF2’s intrinsic energy preferences being unmasked after clustering has removed signal for the other state. Finally, we use AF2 to identify a minimal set of two point mutations predicted to switch KaiB between its two states.

We further hypothesized that our clustering method might be able to detect alternate conformations in protein families for which no alternate structures are known. We applied our method to an existing database of MSAs associated with crystal structures^30^ to aim to detect novel conformational states in known protein families. We describe here one candidate from our screen, the oxidoreductase DsbE. Like the known fold-switcher KaiB, DsbE is predicted to occupy both a thioredoxin-like fold and a novel fold. Our results demonstrate that in the oncoming age of AF2-enabled structural biology, related sequences for any given protein target might harbor signal for more than one biologically-relevant structure, and that deep learning methods can be used to detect and analyze these multiple conformational states.

## Results

### Clustering input MSA sequences by similarity results in AF2 predictions of both KaiB states

We started our investigation with a contradiction posed by predicting the structure of the metamorphic protein KaiB in AF2. KaiB is a circadian protein found in cyanobacteria^8^ that adopts two conformations with distinct secondary structures as part of its function: during the day, it primarily adopts the “ground state” conformation which has a secondary structure of βαββααβ not found elsewhere in the PDB (**Figure 1A**, PDB: 2QKE). At night, it binds KaiC in a “fold-switch” (FS) conformation, which has a thioredoxin-like secondary structure (βαβαββα) (**Figure 1B**, PDB: 5JYT). The thermodynamically favored state for KaiB from *Thermosynechococcus elongatus* (KaiB^TE^) is the ground state; the FS structure was first solved in complex with KaiC^31^, and was solved for the isolated KaiB by introducing stabilizing mutations to this variant^31^. However, AF2 in ColabFold^32^ predicts the thermodynamically unfavored FS state for KaiB^TE^ (**Figure 1C**).

**Figure 1:**
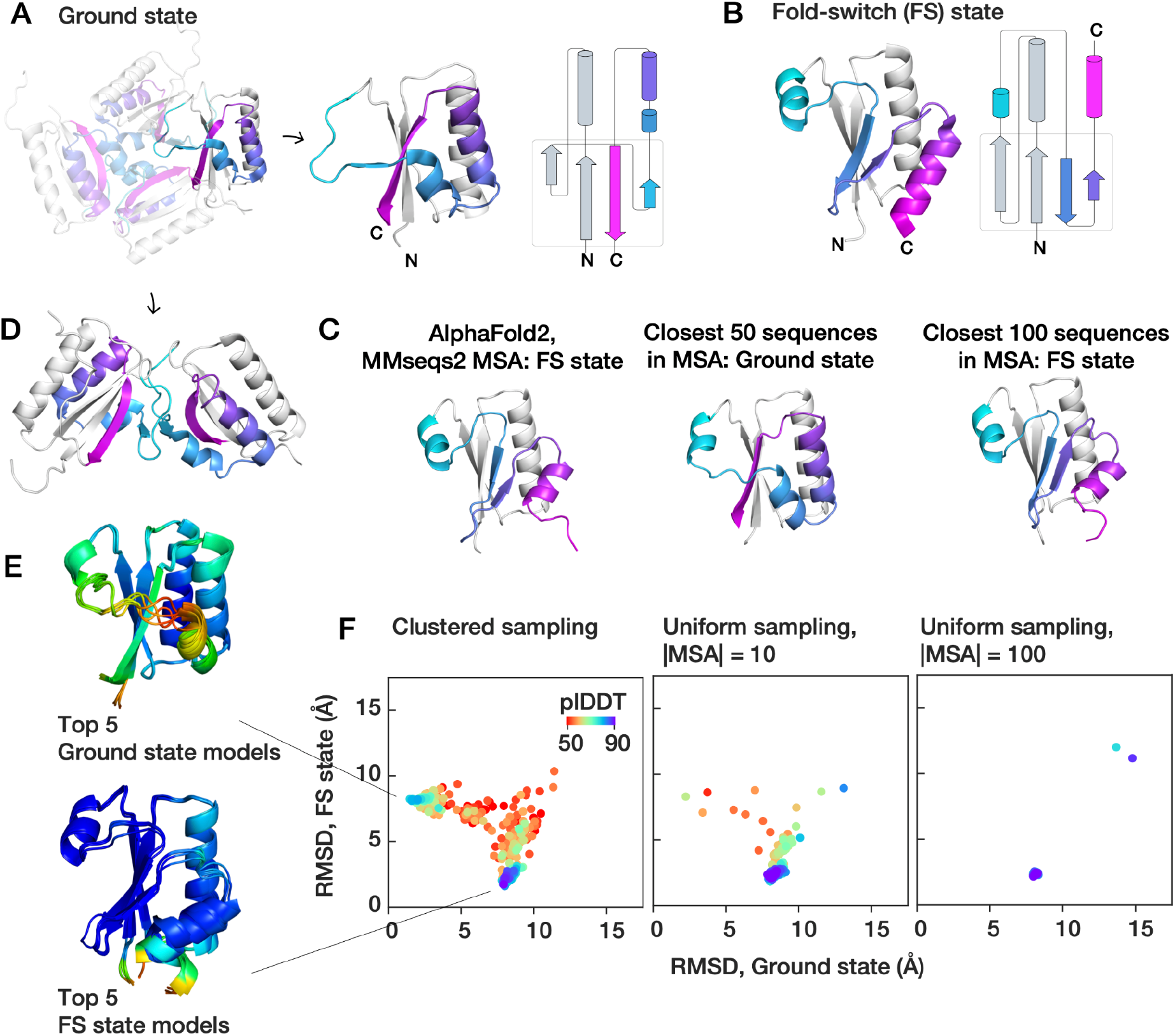
AF2 predictions from MSA clusters for the fold-switching protein KaiB return both known structures. Crystal structures of KaiB of (A) the “ground” state from T. elongatus (KaiB^TE^) (PDB: 2QKE, left: tetramer, right: monomer) and (B) its “fold-switch” (FS) state (PDB: 5JYT). C) The default ColabFold prediction of KaiB^TE^ returns the FS state. Using only the closest 50 sequences by sequence distance returned from the MSA returns the ground state, but the closest 100 returns the fold-switch state. Domain coloring as in Figure 1A. The main discrepancy between the AF2 ground state prediction and the crystal structure is that residues 47-53 are predicted to form a helix; in the crystal structure, these form a dimeric interaction (shown in D). E) Clustering the MSA by sequence distance and predicting for each cluster results in predictions of both states. Top 5 models (ranked by plDDT) within 3 Å RMSD of crystal structures of both states. F) The highest-confidence regions of the entire sampled landscape bear low RMSD to the ground and FS state. In contrast, sampling the MSA uniformly returns only the FS state with high confidence.

We hypothesized that coevolutionary signal within the MSA may be biasing the prediction to the FS state. Interestingly, predicting KaiB using just the 50 sequences from the MSA closest to KaiB^TE^ resulted in a prediction of the ground state; however, predicting KaiB^TE^ using the closest 100 sequences again returned to predicting the FS state (**Figure 1C**). The main discrepancy between the AF2 predictions of the ground state and the tetrameric crystal structure is in the region between 47-53 that form multimeric interactions in the crystal structure (**Figure 1D**).

We thought that the MSA might contain pockets of sequences with signal for either the ground or FS state. Therefore, we clustered the MSA by sequence distance using DBSCAN^33,34^, and ran predictions in AF2 using these clusters as input to AF2. We selected DBSCAN to perform clustering because we reasoned it might offer an automated route to optimizing clustering *a priori* (see Methods, **Figure 2**). From here on we refer to this entire pipeline as “AF-cluster” – generating a MSA with ColabFold, clustering MSA sequences with DBSCAN, and running AF2 predictions for each cluster.

**Figure 2:**
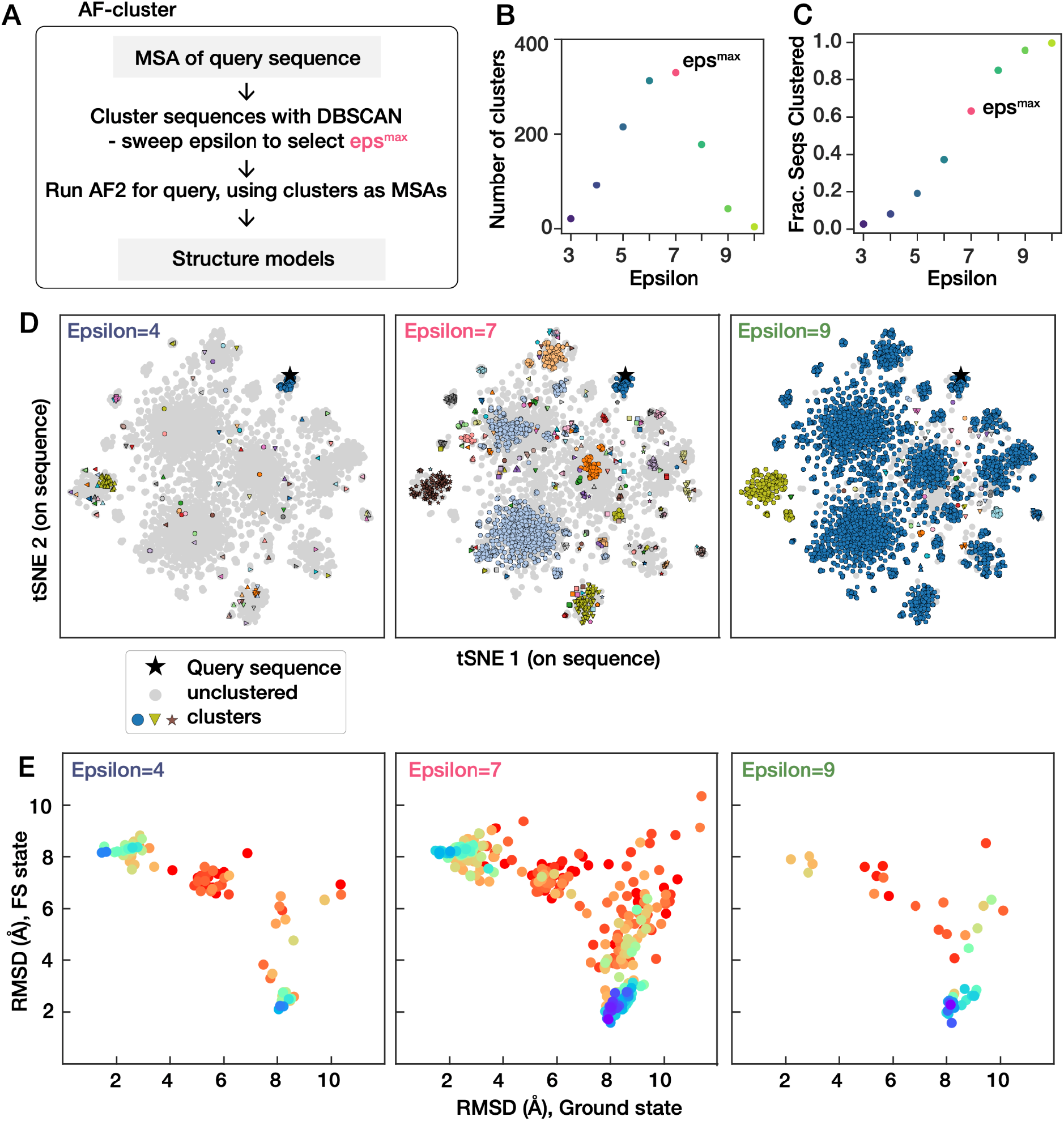
Empirically maximizing information content of clustering using DBSCAN^33^. A) Overview of AF-cluster workflow. B) Varying the parameter epsilon, which controls the maximum allowable distance for points to be in a cluster, results in a peak in the number of clusters DBSCAN identifies for a set of sequences. For eps < eps^max^, fewer sequences are clustered, i.e. more are identified as outliers by the DBSCAN algorithm (C). For eps > eps^max^, more sequences are clustered but fewer clusters are returned as more clusters are joined. (D) Example clusterings of KaiB sequences at different epsilon values. Sequence space is depicted using a tSNE embedding^55^ of the one-hot sequence encoding. (E) Corresponding KaiB landscape of predictions for these values of epsilon.

Strikingly, we found that the AF2 predictions from MSA clusters comprised a distribution of structures, with the highest-scored regions of the distribution corresponding to the ground and FS state. **Figure 1E** depicts the top 5 models within 3 Å of crystal structures for each state, ranked by predicted local distance difference test (plDDT). We compared this subsampling method to predictions from MSAs obtained by uniformly sampling over the MSA at various MSA sizes (**Figure 1F**), analogously to methods used elsewhere to detect conformational changes in GPCRs^18^. We found that for uniformly subsampled MSAs of size 10, one of 500 samples was within 3 Å of the ground state, with lower confidence than the MSA cluster samples (**Figure 1 – Figure supplement 1**). Uniformly subsampled MSAs of size 100 did not sample the ground state at all.

We were curious if the differing signals detected in our MSA clusters by AF2 were also detectable using other deep learning methods, and if this could help us understand how AF2 detected two states. We used the same set of clusters to make predictions with the deep learning model MSA transformer^29^. We found that clusters that predicted the FS state in AF2 also predicted contact maps matching the FS state in MSA Transformer, but clusters predicting the Ground state in AF2 showed little signal in MSA Transformer (see Methods, **Figure 1 – Figure supplement 2**). Some in fact contained more features corresponding to the FS state than the ground state. This suggests there is minimal coevolutionary signal contained in the clusters for the ground state, and that AF2’s prediction of the ground state is instead driven by thermodynamics.

Within our clustering, we noticed conserved sequence motifs enriched within sequence clusters that predicted either the ground or FS state (determined by <3 Å cutoff, **Figure 3A**). It is likely that some of the mutations enriched for either state are due to random evolutionary drift, whereas some play a role in stabilizing or destabilizing one or the other structure. We were curious if we could identify a minimal subset of mutations needed to switch an AF2 prediction from the ground to the FS state (see Methods, **Figure 3 – Figure supplement 1**) to get a better understanding of functional mutations. We used the KaiB^TE^ construct with an MSA comprising the 10 closest sequences from the ColabFold MSA. For each set of mutations tested, we mutated the query sequence as well as every sequence in the MSA. Indeed, we found that the double mutation V68E and I83K were sufficient to switch a prediction of KaiB^TE^ from the ground to the FS state (**Figure 3B**).

**Figure 3:**
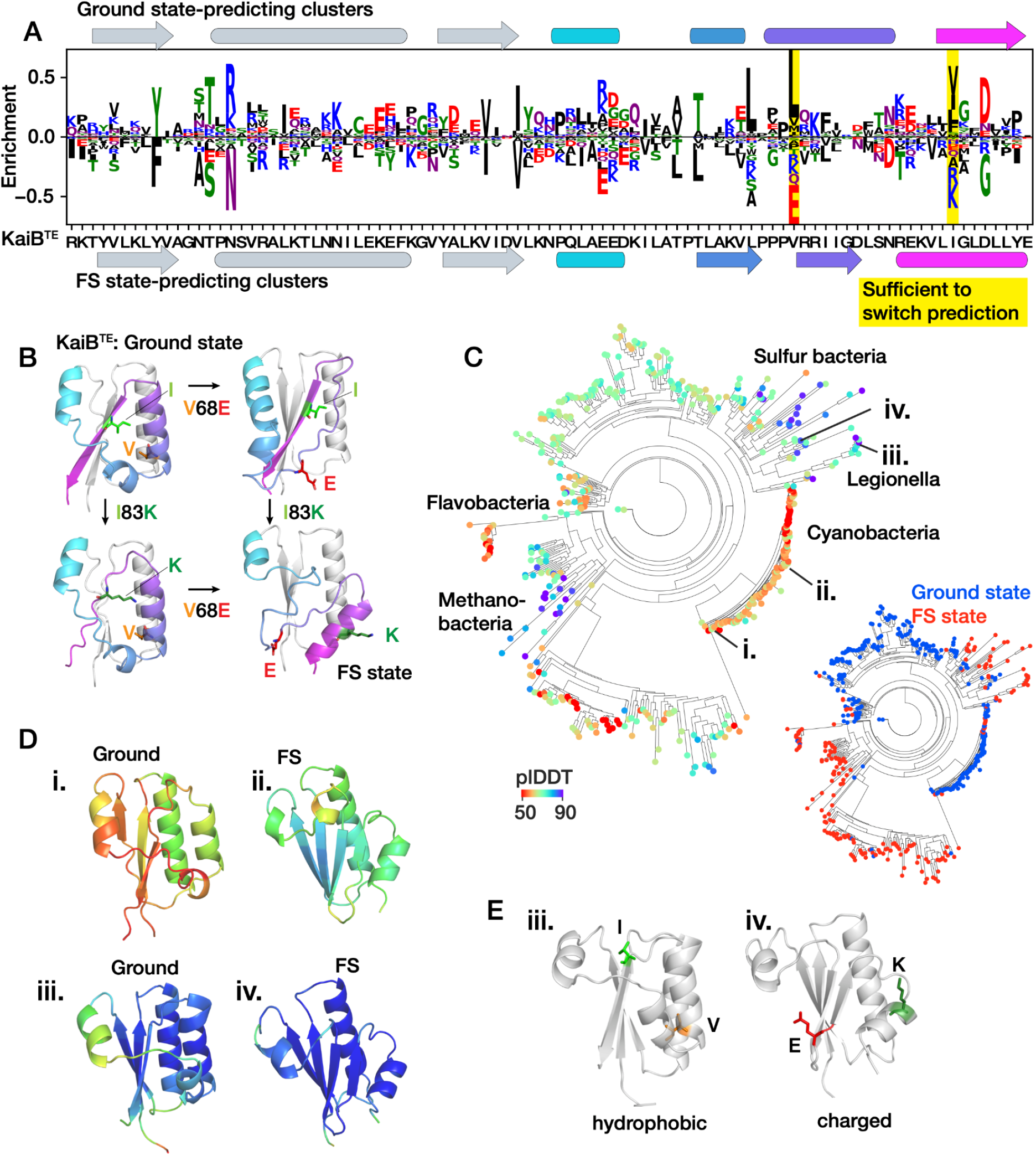
Mutations driving prediction of multiple conformational states in KaiB family. A) Sequence features of clusters predicting ground and FS state. Secondary structure elements of both states are colored as in Figure 1. B) We identified two mutations that are sufficient to switch the structure prediction for KaiB^TE^ from the ground state to the FS state: V68E and I83K, highlighted in yellow in A, and plotted onto the predicted structures for each mutant in B (see **Figure 3 – Figure supplement 1**). C) AF2 predictions for each variant in a phylogenetic tree^35^ using the 10 closest sequences as input MSA. Left: colored by plDDT, right: each node colored by predicted state (blue: ground state, red: FS state). D) Predicted structures of two known fold-switching KaiB variants: *Synechococcus elongatus* (i) and *Thermosynechococcus elongatus* (ii), and two variants with highest plDDT predicted to be stabilized for one fold or the other: *Ectothiorhodospira sp. PHS-1* (iii) and *Legionella pneumophilia* (iv). All structures ccolored by plDDT. E) Two variants in (D) strongly predicted for the ground (iii) and FS state (iv) respectively, colored the same as the point mutations in (B). The variant from *Legionella pneumophilia* contain the exact 2 mutations predicted in B to drive the fold switch.

To better understand the evolutionary relationships of KaiB variants that AF2 predicted to be stabilized for one or the other state, we made structure predictions for 487 KaiB variants in a curated phylogenetic tree^35^ (**Figure 3C**). For each construct, we used only the closest 10 constructs by evolutionary distance as an input MSA to try to best detect local differences in structure predictions. We found that regions of high plDDT for both the ground and FS state were interspersed across the tree. We observed that variants from the cyanobacteria phylum had lower plDDT than other phyla. **Figure 3D** i and ii depict the predicted structures for the two fold-switching species *Synechococcus elongatus and Thermosynechococcus elongatus* (TE). Notably, our previous 10-sequence MSA for KaiB^TE^ from the colabfold MSA predicted the ground state in the previous analysis, but this 10-sequence MSA predicted the FS state, underscoring the variability in predictions for this variant.

In contrast to the cyanobacteria phylum, other phyla contained variants with high-pLDDT predictions for both the ground and FS state. Interestingly, the two mutations identified previously as sufficient to switch AF2 prediction from the ground to the FS state, V68E and I83K (**Figure 3B**), were also present in KaiB variants with high-plDDT FS state predictions from our separate analysis of the KaiB phylogenetic tree (**Figure 3E**). Both mutations change from hydrophobic residues that are buried in the ground state to charged residues that are solvent-exposed in the FS state. Though these mutations need to be experimentally tested, this demonstrates that AF2 had sufficient signal to make a prediction for fold-switching between point mutations and suggests that AF2 predictions could also be applied to understand the evolution of fold-switching evolutionary paths for metamorphic proteins.

### AF-cluster predicts monomeric alternate states in other metamorphic proteins

We next tested AF-cluster on five other experimentally verified fold-switching proteins: the *E. coli* transcription and translation factor RfaH, the human cell cycle checkpoint Mad2, the selecase metallopeptidase enzyme from *M. janaschii*, the human cytokine lymphotactin, and the human chloride channel CLIC1. In RfaH, the C-terminal domain (CTD) interconverts between an α-helix bundle and a β-barrel through binding to functional partners^10^. In the autoinhibited state, the α-helix bundle of the CTD interacts with the NTD. In the active state, the CTD unbinds and forms the β-barrel (**Figure 4A**)^9,10^. Predicting the structure of RfaH with the complete MSA from ColabFold returned a structure that largely matched the autoinhibited state (**Figure 4 – Figure Supplement 1A**) apart from the first helical turn in the CTD being predicted as disordered. We note that the B-factors in the crystal structure for this region are highest (**Figure 4 – Figure Supplement 1B**). The active state was not predicted. In contrast, AF-cluster predicted both the autoinhibited and the active state (**Figure 4B**). Notably, the average plDDT for the top 5 models for each state (84.2 for the active state, 73.9 for the autoinhibited) was higher than the plDDT of the autoinhibited state by the complete MSA (pLDDT of 68.6), suggesting that clustering resulted in deconvolving conflicting coevolutionary signals.

**Figure 4:**
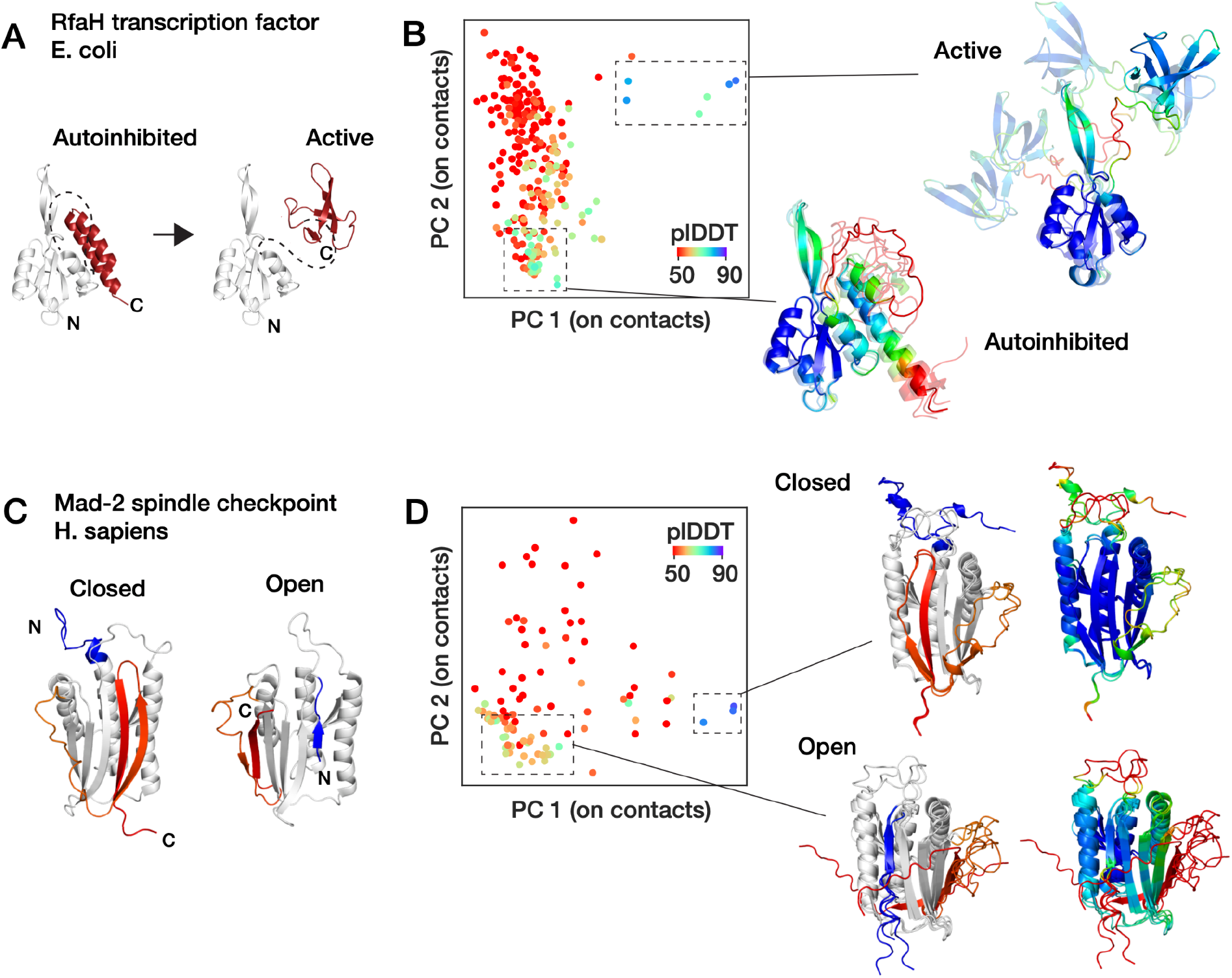
AF-cluster predicts fold-switching for the proteins RfaH and Mad2. A) Scheme of foldswitching in the RfaH transcription factor in *E. coli*. In RfaH’s autoinhibited state, the C-terminal domain (CTD) forms an alpha-helix bundle. In the active state, the CTD unbinds and forms a β-sheet that is homologous to the transcription factor NusG. D) AF-cluster returns structure models that include both the autoinhibited and the active state, both with higher plDDT scores than the model of the autoinhibited state in (B). E) Two states of the Mad-2 spindle checkpoint in humans with the fold-switching portions colored. F) Both Mad-2 states are predicted by AF-cluster.

Mad2 has two topologically distinct monomeric structures that are in equilibrium under physiological conditions^15^. These are termed the open and closed states (often referred to as O-Mad2 and C-Mad2). The closed state binds Cdc20 as part of Mad2’s function as a cell cycle checkpoint^13^. In the closed state, the C-terminal beta-hairpin rearrange into a new β-hairpin that binds to a completely different site, displacing the original N-terminal β-strand^16^ (**Figure 4C**). We found AF-cluster was again capable of predicting models for both of Mad2’s conformational states (**Figure 4D**).

RfaH and Mad2 both interconvert between two distinct monomeric forms. However, selecase, lymphotactin, and CLIC1 both interconvert between a monomeric and an oligomeric state (**Figure 4 – Figure supplement 2**). AF-cluster was unable to predict the oligomeric state for selecase, lymphotactin, and CLIC1. The selecase protein is a metallopeptidase from M. janaschii first reported by Lopez-Pelegrin et al^11^. It reversibly interconverts between an active monomeric form and inactive dimers and tetramers. Lymphotactin is a human cytokine that adopts a cytokine-like fold but was found to adopt an all-β-sheet dimer via NMR at higher temperature and in the absence of salt^12^. CLIC1 is an ion channel with a redox-enabled conformational switch. In the reduced state, it adopts a monomeric state with a N-terminal βαβαβ fold. Upon being oxidized, it forms a dimer, and its N-terminus adopts a ααα fold. This fold is stabilized by a disulfide bond between two of the *α*-helices within the monomer that forms upon oxidation^36^. All these proteins pose starting points for future improvements to AF-cluster.

### Screening with AF-cluster predicts fold-switching in another thioredoxin-like protein

Despite its limitation in only predicting monomeric alternate states, we next asked if AF-cluster could detect novel putative alternate states in protein families without known fold-switching (**Figure 5A**). As a starting point, we selected 628 proteins with length 48-150 amino acids from a database of MSAs associated with crystal structures^30^ (see Methods). After clustering the MSAs using DBSCAN^33^, we generated AF2 predictions for 10 randomly-chosen clusters from each family and compared the plDDT to the RMSD from the reference structure (RMSD^ref^). For most of the protein families screened, an increase in RMSD^ref^ corresponded to a decrease in plDDT (**Figure 5B**). However, a handful of proteins in this preliminary screen returned models with high RMSD^ref^ and high plDDT, indicating a predicted structure with high dissimilarity to the original structure as well as high confidence from AF2. For these proteins, we generated AF2 predictions for all generated clusters from the MSA.

**Figure 5:**
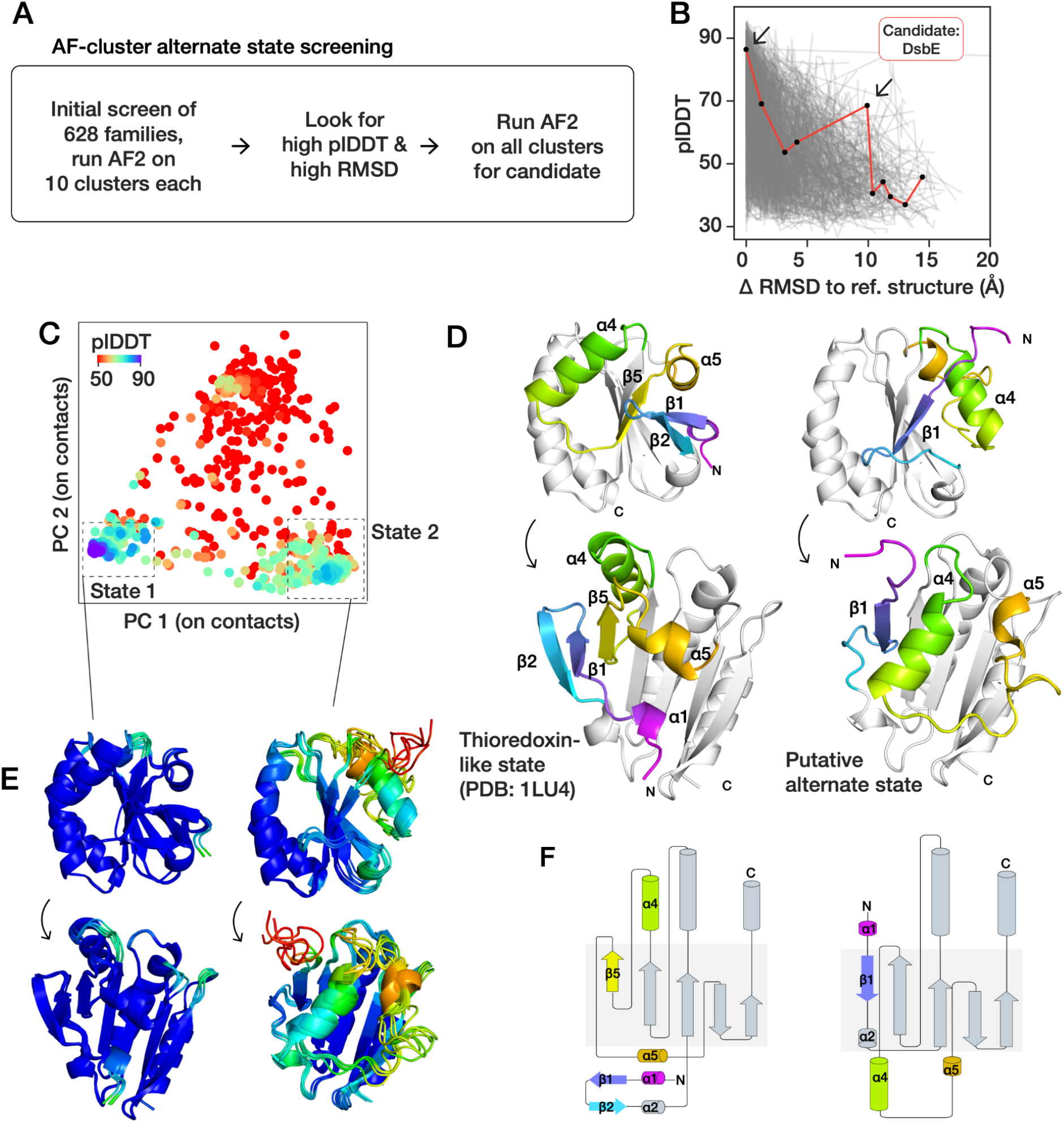
Screening for fold-switching predicts a putative alternate fold for *M. tuberculosis* oxidoreductase DsbE (Mtb DsbE). A) Strategy for detecting novel predicted alternate folds. We screened 628 families with more than 1000 sequences in their MSA and residue length 48-150 from ref.^30^. After clustering, we ran AF2 predictions from 10 randomly-selected clusters from each. B) We selected candidates for further sampling by looking for outlier predictions with high RMSD to the reference structure and high plDDT. C) Sampled models for candidate Mtb DsbE, visualized with a principal component analysis on closest heavy-atom contacts. Two states with higher plDDT than background are observed. D) Left: Crystal structure of Mtb DsbE (PDB: 1LU4), which corresponds to state 1 in the sampled landscape, has a canonical thioredoxin fold. In the putative alternate state 2, strand β5 replaces β1 in the 5-strand β-sheet. Helix α4 shifts to the other side of the β-sheet and helix α5 is displaced. E) Top 5 models by plDDT for the thioredoxin-like state (left) and the putative alternate state (right), colored by plDDT per residue. F) Secondary structure diagrams for DsbE thioredoxin-like state in crystal structure 1LU4 and putative alternate state.

Results for one of these fold-switching candidates, the oxidoreductase DsbE from *M. tuberculosis* (Mtb DsbE), is described here. Mtb DsbE is an extracellular single domain enzyme that ensures correct folding of several cell-wall and extracellular protein substrates in M. tuberculosis by catalyzing disulfide oxidoreduction^37–39^. **Figure 5C** depicts all the AF-cluster models for Mtb DsbE, visualized by principal component analysis (PCA) on the set of closest heavy-atom contact distances. Two prominent states are observed that correspond to the largest-sized MSA clusters (**Figure 5 – Figure Supplement 1A**), and both of which have plDDT values statistically significantly higher than the rest of the set (**Figure 5 – Figure Supplement 1B**). One state corresponds to the canonical known thioredoxin-like conformation of DsbE^38^, whereas the other state corresponds to an unknown conformation with a different secondary structure layout (**Figure 5D-F**). In the second state, the strand β5 is switched with β1 in the β-sheet. The α-helix α4 is displaced to the opposite side of the β-sheet, and α5 is rotated. Mtb DsbE is a member of a superfamily of enzymes with diverse functions that all share the same thioredoxin fold with a conserved CxxC active site with a disulfide bond. Models for the alternate state demonstrate the same active site orientation at residues C36-C39 (**Figure 5 – Figure Supplement 1C**). We screened for homologous 3D structures for the alternate state in the PDB using DALI^40^. The top 30 hits resembled the helical orientations of the known thioredoxin-like fold of DsbE (**Figure 5 – Figure Supplement 2**).

We next tested if any structure homologues to DsbE also predicted alternate conformations. We identified 6 structure homologues (**Figure 6A**) using 3D-Blast^41^, which ranged in sequence similarity to DsbE from 22% to 13%, to test for homologous alternate states. Using AF-cluster, 4 of the 6 sampled an analogous alternate fold (**Figure 6B-D**): the soluble domains of ResA from *B. subtilis*, the periplasmic N-terminal soluble domain of DsbD from *E. coli*, TlpA from *B. japonicum*, and Sco1 from humans. For all except human Sco1, the alternate state was also scored higher than the background. This demonstrates a range of confidence in predictions and allows for experimentally testable hypotheses going forward.

**Figure 6:**
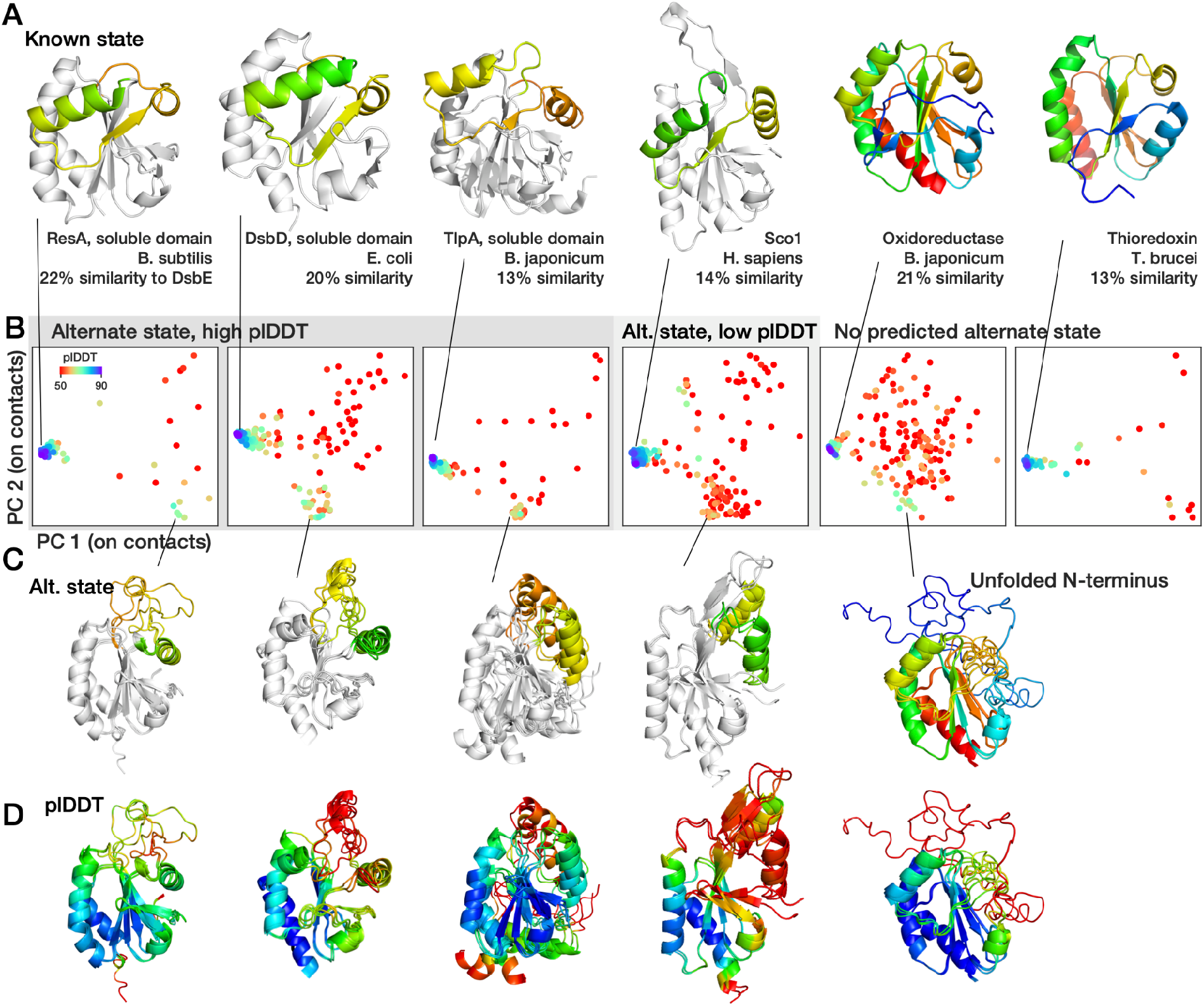
An analogous fold-switch state is predicted for some DsbE structure homologues. A) Crystal structures of 6 proteins with high structure homology to Mtb DsbE. PDB codes from left to right: 1ST9, 1Z5Y, 1JFU, 1WP0, 1KNG, 1R26. Those with predicted fold-switching (first 4) are colored in the region that fold switch with grey in regions that the same. B) Conformational landscapes of AF-cluster predictions for these 6 proteins range from predicting an analogous alternate state with high confidence to showing no prediction of the alternate state. C) Predicted alternate structures. The alternate structure for the oxidoreductase from *B. japonicum* is not the conserved putative alternate state, but instead an unfolded N-terminus. D) Alternate structures in (C), colored by plDDT. The alternate structure for *H. sapiens* does not have a plDDT above background.

## Discussion

AlphaFold2 (AF2) has revolutionized prediction of single structures^42^, but devising methods to predict structures of multiple conformational states would significantly advance our understanding of protein function at atomic resolution. We demonstrate that simply clustering input sequences from MSAs of metamorphic proteins enables AF2 to both sample multiple conformations and score them with high plDDT. We use our method to identify alternate states in protein families without known fold-switching, and speculate that there may be more previously uncharacterized functional states present that this method could uncover. The AlphaFold protein structure prediction database^43^ contained 214 million predictions of single structures as of September 2022. If Porter and Looger’s estimate ^17^ that 0.5-4% of all proteins contain fold-switching domains is accurate, this would correspond to roughly 1-8 million fold-switching proteins present. However, our findings demonstrate that making a single structure prediction from a given MSA can mask signals of alternate states.

We used the protein language model MSA Transformer^29^ in conjunction with AF2 to better understand the two sets of signals for the metamorphic protein KaiB. The subsampled MSA clusters that predicted the FS state in AF2 also predicted the FS state in MSA Transformer, but we found a discrepancy between the two methods in predicting the ground state. Clusters that predicted the ground state in AF2 had little signal for the in MSA Transformer contact map predictions. We speculate that AF2 predicted the FS state through detecting coevolutionary information, but that the ground state prediction came from intrinsic AF2 “thermodynamics” after clustering removed conflicting coevolutionary signal for the FS state. This suggests that coevolutionary signal input via an MSA and intrinsic preferences of AF2 do not always favor the same conformational state, and that these discrepancies can be leveraged to uncover multiple states.

Furthermore, we identified a minimal set of 2 mutations sufficient to switch the prediction of KaiB^TE^ from the ground to FS state and found that portions of a curated phylogenetic tree for KaiB contained high-confidence AF2 predictions for both states which contained these mutations. We hypothesize those KaiB variants are thermodynamically stabilized for those states, but experimental testing is needed. It would not be surprising that the KaiB family would contain pockets of constructs that are stabilized for one or the other, as has been found for the fold-switchers RfaH^44^ and lymphotactin^45^, as well as non-fold-switching proteins like the Cro repressor family^46^. This demonstration that point mutations can alter AF2 predictions from a shallow MSA comprising 10 closely-related sequences suggests that other works demonstrating AF2 is incapable of predicting changes in thermodynamic stability due to point mutations^47^ might be revisited. Disease-causing point mutations are often due to population changes of protein substates^48,49^, and thus there is significant need for methods better able to predict the effects on free energy and conformation of point mutations.

By using AF-cluster to screen protein families not known to fold-switch into alternate states, we identified a putative alternate state for the oxidoreductase DsbE. The thioredoxin superfamily containing DsbE is a ubiquitous set of enzymes known for their promiscuous catalytic activity, being able to reduce, oxidize, and isomerize disulfide bonds^50^. Theoretical work suggests that conformational change is the most parsimonious explanation of the evolution of promiscuous activity in the thioredoxin family^51^. To the best of our knowledge, the one example of secondary structure rearrangement in an oxidoreductase is suggestion of a secondary structure rearrangement in a related oxidoreductase protein in the thermophile *Pyrococcus furiosus* to explain melting data^52^. No structure model for the alternate state was proposed. AF-cluster predicts a high-plDDT rearrangement of two of the 8 β-strands repacking (**Figure 6 – Figure supplement 1**). Given that known metamorphic proteins switch folds through cellular stimuli, it may in general be difficult to validate novel folds identified through computational methods if the stimulus – whether pH, redox reaction, a binding partner – is unknown.

Further study is ongoing in what types of conformational changes AF-cluster and other methods based on altering input MSAs can predict. Because previous works have identified coevolutionary couplings corresponding to multiple states of domain-based conformational changes, we speculate that clustering-based MSA preprocessing methods will offer improvements over existing methods^18^, and importantly, insights in the evolution of multiple conformational states. However, conformational changes not present in coevolutionary signal may require alternate methods. All methods also need to be evaluated and improved in their ability to sample and score in accordance with the system’s underlying Boltzmann distribution. As protein sequencing data continues to increase, computational methods for characterizing and identifying conformational motions will likely provide increasing insight into protein folding, allostery, and function.

## Acknowledgments

We thank Hannes Ludewig, Ricardo Padua, Marc Hömberger, Renee Otten, and other members of the Kern lab for helpful discussions and feedback. AlphaFold2 calculations were run on the Harvard Medical School O2 cluster. H.K.W-S. acknowledges funding from the Jane Coffin Childs foundation. This work was supported by the Howard Hughes Medical Institute (HHMI) to D.K.

## Author Contributions

H.K.W-S. and D.K. conceived the project. H.K.W-S., S.O., L.C., and D.K. designed experiments. H.K.W-S. performed all calculations and analysis. H.K.W-S. and D.K. wrote the paper. H.K.W-S., S.O., L.C., and D.K. commented on the manuscript and contributed to data interpretation.

## Methods

### MSA generation

Multiple sequence alignments (MSAs) were generated using the MMseqs2^53^-based routine implemented in ColabFold^32^. In brief, the ColabFold MSA generation routine searches the query sequence in three iterations against consensus sequences from the UniRef30 database^54^. Hits are accepted with an E-value lower than 0.1. For each hit, its respective UniRef100 cluster member is realigned to the profile generated in the last iterative search, filtered such that no cluster has higher max sequence identity than 95%, and added to the MSA. Additionally, in the last round of MSA construction, sequences are filtered to keep the 3000 most diverse sequences in the sequence identity buckets [0.0–0.2], (0.2–0.4], (0.4–0.6], (0.6–0.8] and (0.8–1.0].^32^ Prior to clustering, we removed sequences from the MSA containing more than 25% gaps.

### Clustering

We found that our method for parameter selection in DBSCAN^33^ empirically optimized predicting KaiB’s two states from the MSA with no prior information about the KaiB landscape. A schematic of the AF-cluster method is depicted in **Figure 2A**. An optimal clustering to identify signal of multiple states needs to balance two size effects: if clusters are too small, they may contain insufficient signal to capture any state. However, if clusters are too large, they may dilute signal from some states, an extreme case of this being how KaiB predicted with its entire MSA resulted in only the fold-switch state. In brief, DBSCAN^33^ clusters datapoints by identifying “core” density regions where at least *k* points fall within distance *epsilon* from one another. Points farther than epsilon from points in core density regions are excluded as noise. Clustering the KaiB MSA with varying epsilon values resulted in a peak in the number oτ clusters returned (**Figure 2B**). We termed the epsilon corresponding to this peak eps ^max^. For eps < eps^max^, the number of clusters is lower because more sequences are left unclustered as outliers (**Figure 2C**). For eps > eps^max^, more sequences are clustered, so the number of clusters is decreasing because clusters are merged.

We investigated the effect of varying epsilon on resulting AF2 predictions for the protein KaiB. **Figure 2D** depicts clusters in sequence space (represented with tSNE^55^ on sequence one-hot encoding), and **Figure 2E** depicts the structure landscape of these clusters. Epsilon=7 was the value used for the results described in the main text. We found that predictions at eps^max^ balanced the number of models returned for each state (**Figure 2 – Figure Supplement 1A**) and their resulting plDDT (**Figure 2 – Figure Supplement 1B**). For the preliminary scan of 628 protein families, this sweep on epsilon was performed on a randomly-selected 25% of the MSA to accelerate computation. Epsilon was varied between 3 and 20 with step size 0.5.

### MSA Transformer predictions

We made predictions for DBSCAN-generated KaiB clusters in the model MSA Transformer^29^ using default settings. For clusters with more than 128 sequences, the “greedy subsampling” routine was used to select sequences.

We scored predicted contact maps to the KaiB ground and FS state in the following way. If 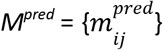 is a NxN array representing the predicted contact map from MSA Transformer, then the Ground state score is given as

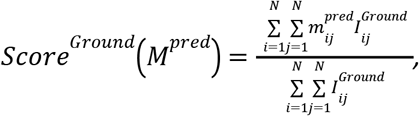

where 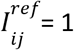 if i and j are paired in the Ground state and not the FS state, and 0 otherwise. The FS state score follows analogously. The contact maps used for this scoring are depicted in **Figure 1 – Figure Supplement 2A**. Every cluster was thus assigned a corresponding “Ground state score” and “FS state score” reflecting its similarity to both states.

We found that the FS state score negatively correlated with the RMSD to the FS state of AF2 (RMSD^FS^) (Spearman R = −0.55, **Figure 1 – Figure Supplement 2B**), indicating that both methods – AF2 and MSA Transformer -- agreed on the amount of signal for the FS state that they detected within a cluster. Contact map features corresponding to the FS state can be identified in the top clusters ranked both by RMSD^FS^ and the FS state score (**Figure 1 – Figure Supplement 2C**). However, the ground state scores for all clusters were lower than the FS state scores. The ground state score and RMSD^Ground^ correlated less (Spearman R=-0.36). The top clusters by ground state score had contact maps corresponding to the antiparallel beta strand that is unique to the ground state (outlined in blue in **Figure 1 – Figure Supplement 2A,C**), but the top clusters by RMSD^Ground^ did not have noticeable ground state features; instead, some exhibit FS state features (outlined in orange, magenta, and red, **Figure 1 – Figure Supplement 2C**). This indicates that the clusters that predicted the Ground state in AF2 did not have coevolutionary signal for the ground state that MSA Transformer recognized, and suggests that AF2 was able to predict the ground state through its learned energy function.

### Data selection for fold-switching screening

Protein families were selected from a database previously developed to query the origins of spatially distant coevolutionary contacts^30^, and last updated in 2017. The database consisted of nonredundant proteins with associated X-ray structures with resolution < 2 Å. The MSAs were originally constructed using HHblits^56^ run against the UniProt database and filtered to exclude sequences with high similarity^30^. Though the database originally contained 9,846 proteins, for this preliminary work we only selected proteins with sequence length between 52 and 150 residues and with more than 1000 sequences in the alignment, which totaled 628 proteins.

### Analysis

Root-mean-squared deviation (RMSD) for structure models was calculated in MDtraj^57^. Principal component analysis and t-SNE^55^ were calculated using Scikit-learn^58^. Spearman correlations and t-tests were calculated using Scipy^59^. Protein structures were visualized in PyMOL^60^.

### Data availability

Scripts for running AF-cluster, AF2, MSA transformer, and data corresponding to structure models and analysis presented here are available at www.github.com/HWaymentSteele/AFCluster.

## Supplemental Figures

**Figure 1 – Figure supplement 1.**
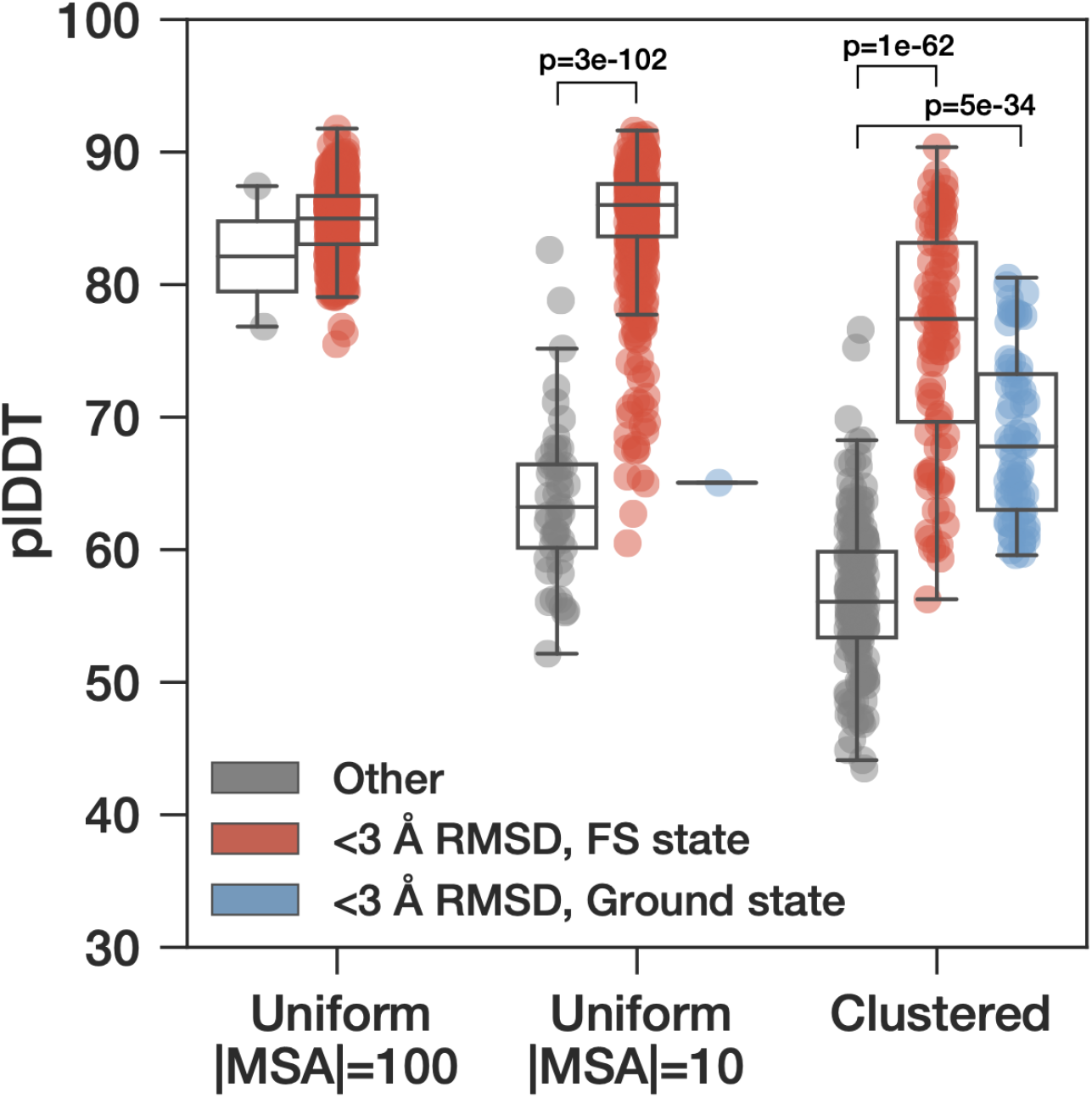
plDDT values of sampled models for KaiB^TE^ from 3 subsampling methods. The plDDT values of models within 3 Å RMSD of the ground- and FS-state from the clustered sampling method are statistically significantly higher than the rest of the models. Box plots depict median and 25/75% interquartile range, whiskers = 1.5*interquartile range. P-values for sample comparisons with p < 0.05 indicated, calculated via a two-sided test for the null hypothesis that 2 independent samples have identical mean values.

**Figure 1 – Figure supplement 2:**
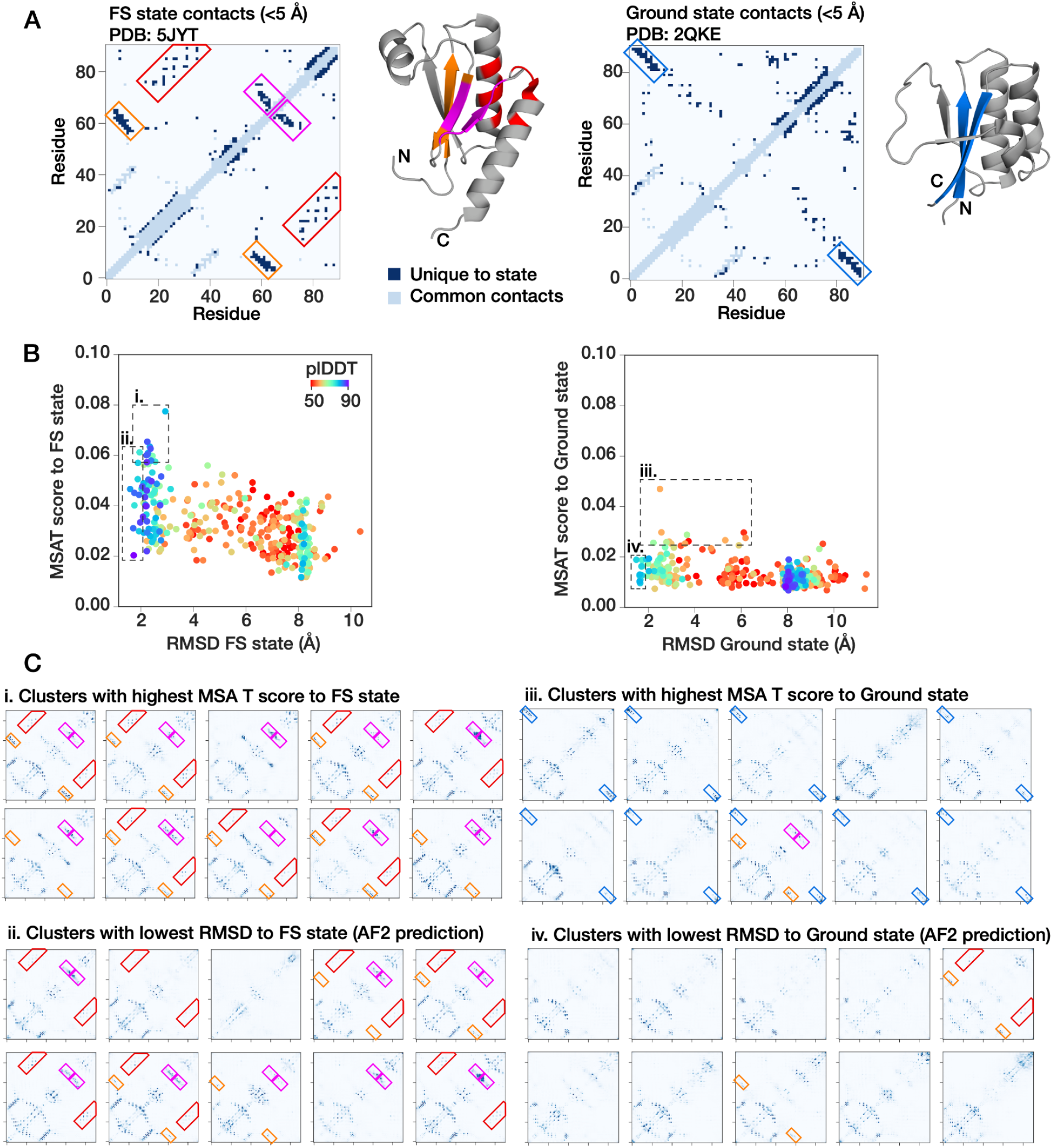
MSA Transformer also detects FS state, but ground state signal is weaker. A) Contacts under 5 Å that correspond uniquely to the FS state (left) or ground state (right). B) FS state scores correlate to the AF2 prediction RMSD to the FS state (Spearman R = −0.55), but ground state scores have less range and weaker correlation (Spearman R = −0.36), colored by plDDT of AF2 prediction. C) Contact maps predicted by MSA Transformer of example clusters optimizing either the MSA Transformer score or AF2 RMSD for both states. Clusters that best predict the FS state in MSA Transformer (i) and AF2 (ii) show features corresponding to beta-strands (orange, magenta) and the helix-helix interaction (red) boxed in (A). Clusters with the highest score to the ground state from MSA transformer (iii) show signal for the beta-strand interaction (blue) unique to the ground state. However, clusters that predict the ground state in AF2 (iv) show minimal features corresponding to the ground state in contact maps predicted by MSA Transformer. Some instead contain FS state features (features boxed in orange, magenta, red with same feature coloring as in A).

**Figure 2 – Figure supplement 1.**
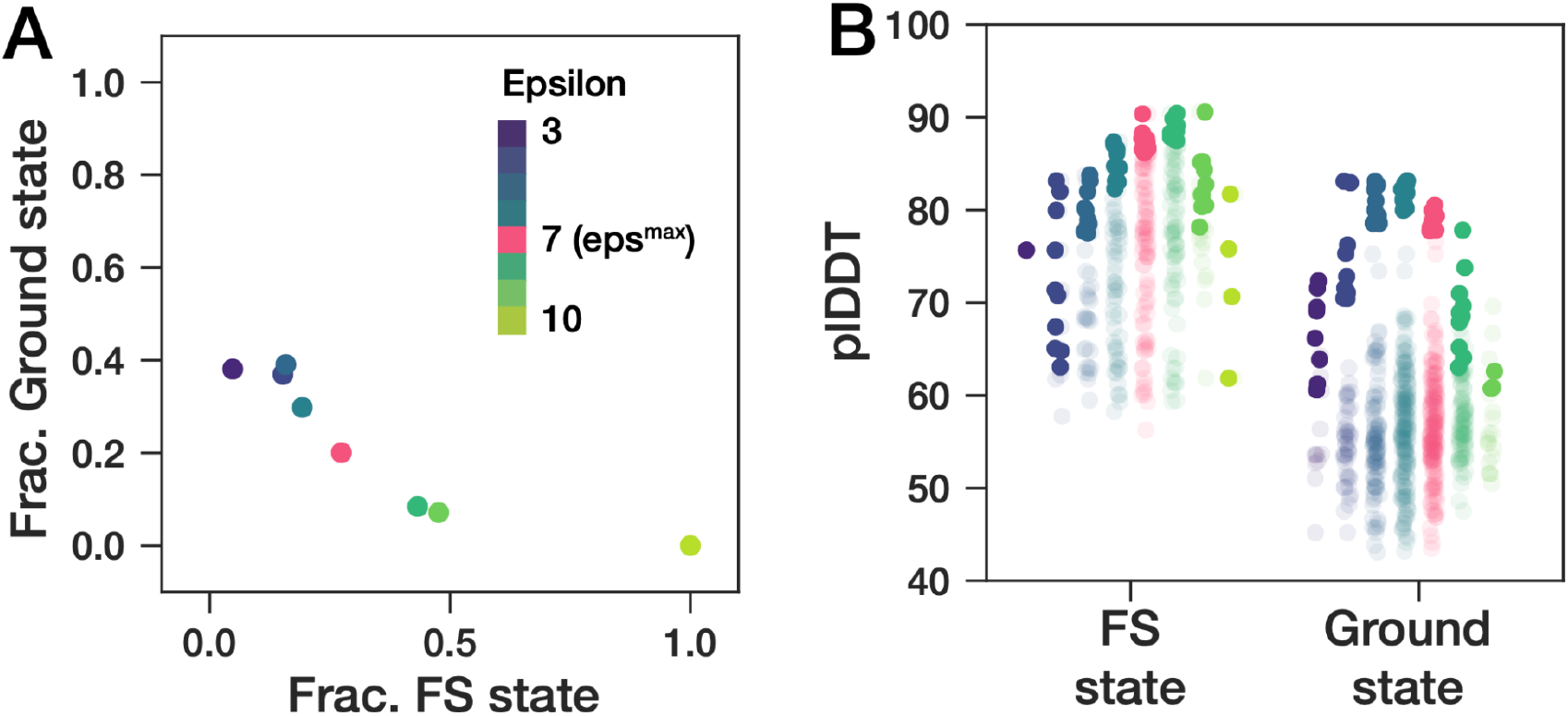
Properties of AF2-predicted structures at different epsilon values. A) As epsilon increases, a higher fraction of the total models correspond to the FS state. (B) plDDT of the two KaiB states. Clustering at eps^max^ returns a mean plDDT that is not statistically significantly different than the eps values for each state that return the highest mean plDDT. The top 10 models are shaded solid, the rest of the models are semi-transparent.

**Figure 3 – Figure Supplement 1:**
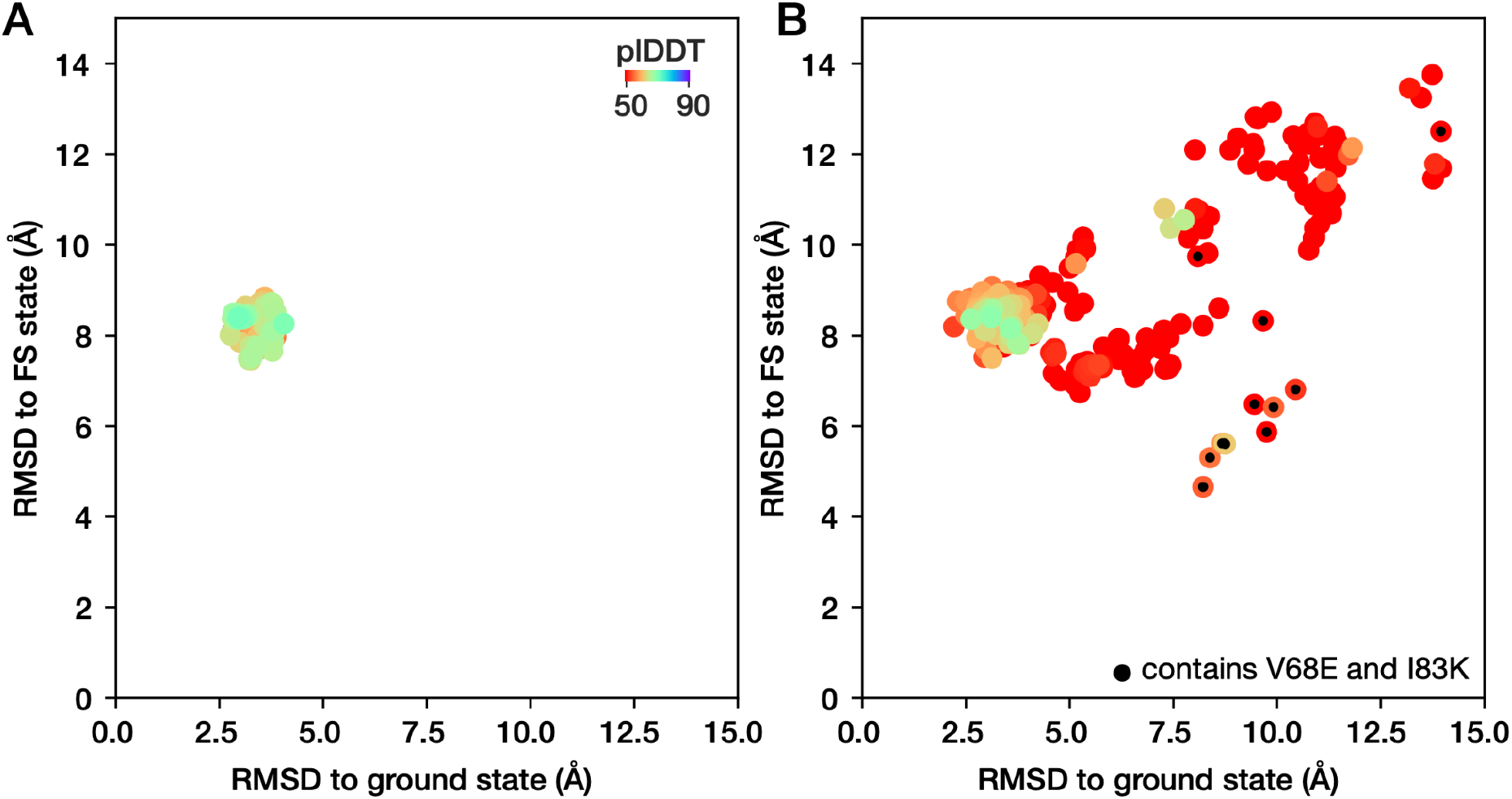
Identifying minimal mutations needed to switch structure prediction for KaiB^TE^. RMSD of the predicted structures for all mutations to (A) the ground state or (B) the FS state. The double mutation V68E and I83K is the only one to cause predictions for KaiB^TE^ to switch states, indicated by a decrease in RMSD to the FS state.

**Figure 4 – Figure Supplement 1.**
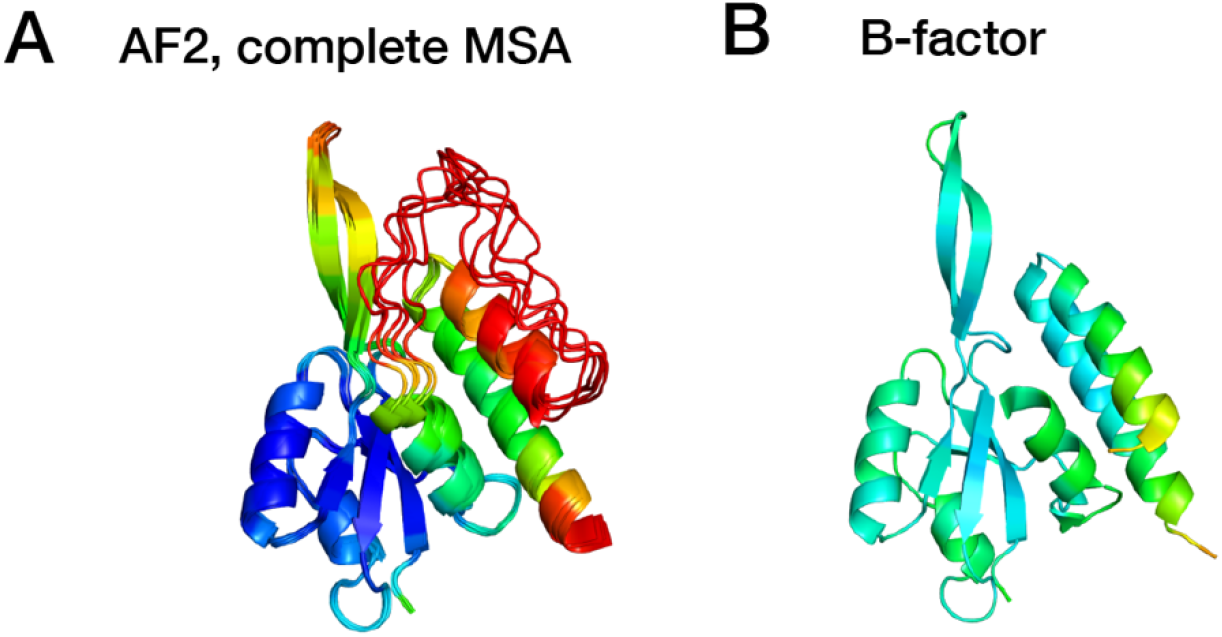
A) Predicting the structure of RfaH in AF2 with the complete MSA from ColabFold/mMseqs2 returns the autoinhibited state with a mean plDDT of 68.6 (note low confidence in the first alpha-helix of the CTD.) B) B-factors of PDB model 5OND, indicating that the last helical turn of the second to last helix has high B-factors.

**Figure 4 – Figure Supplement 2:**
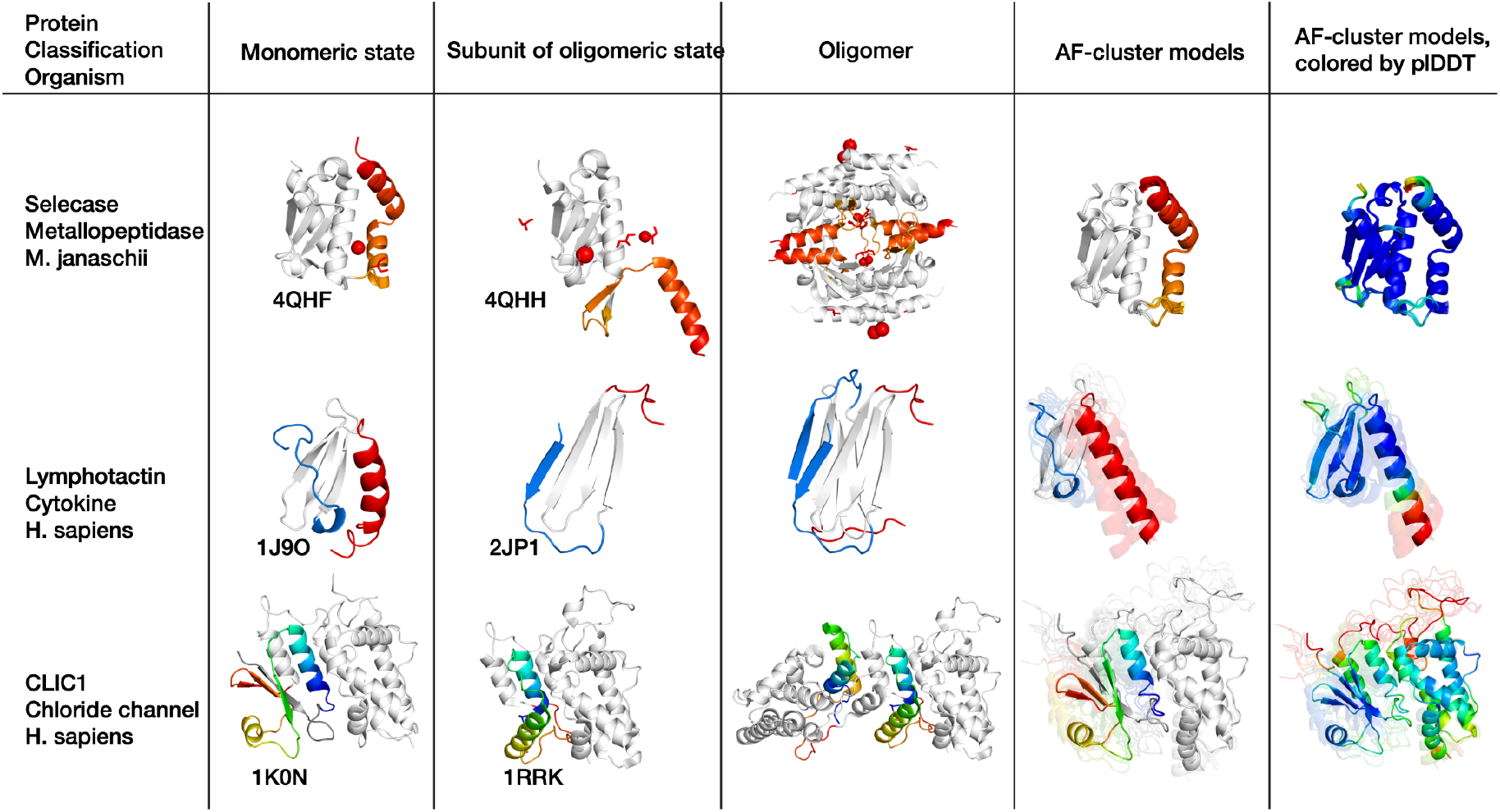
AF-cluster only predicts the monomeric state for proteins that switch between monomeric and oligomeric states.

**Figure 5 – Figure Supplement 1.**
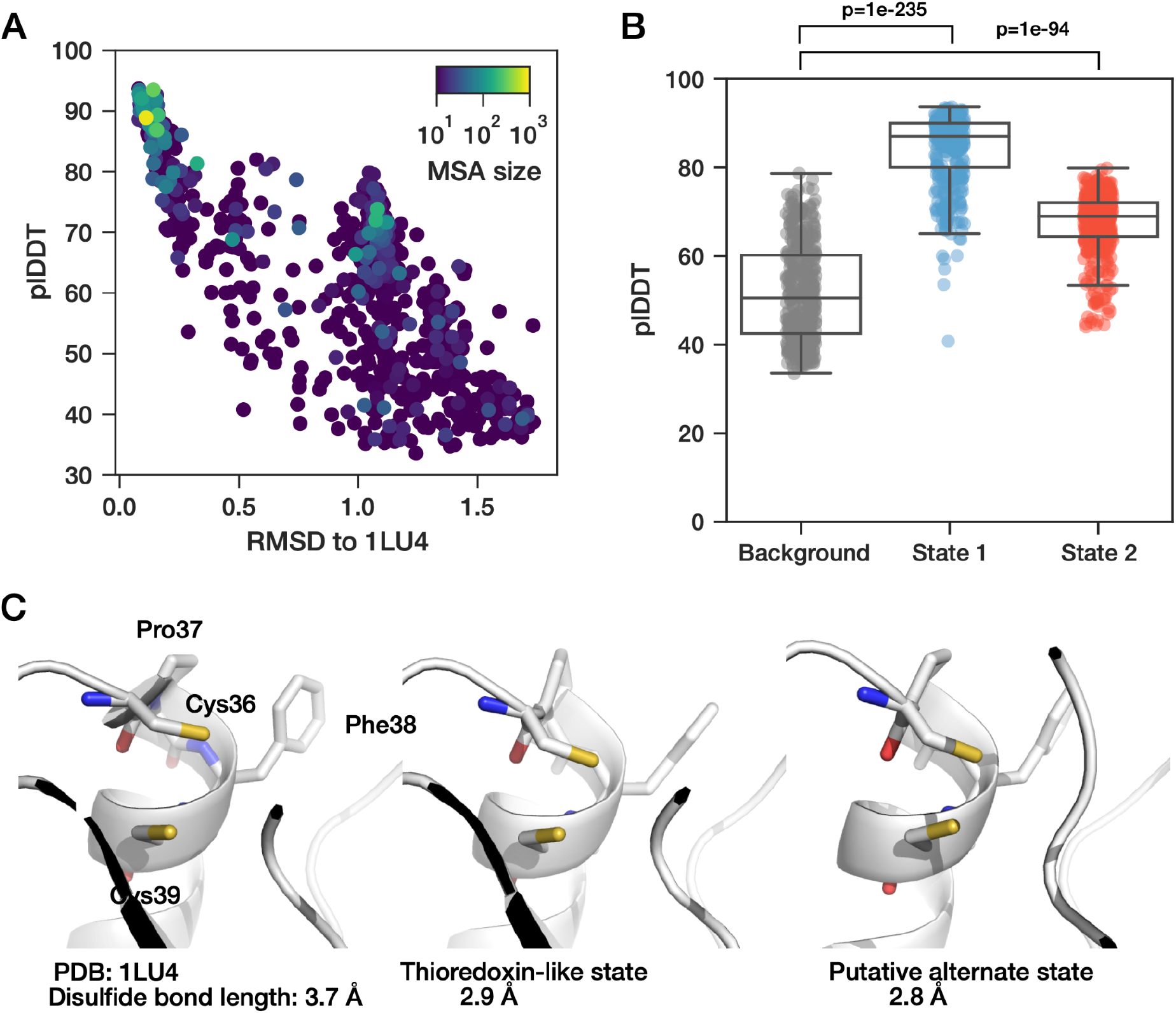
**(A)** plDDT vs. RMSD for increased sampling on oxidoreductase DsbE from *M. tuberculosis*. Each prediction colored by MSA size. (B) plDDT values for state 1, corresponding to the known thioredoxin-like state, and an alternate unknown state are significantly higher than background. Box plots depict median and 25/75% interquartile range, whiskers = 1.5*interquartile range. P-values for sample comparisons with p < 0.05 indicated, calculated via a two-sided test for the null hypothesis that 2 independent samples have identical mean values. (C) The conserved CxxC catalytic domain is unchanged between its conformation in the crystal structure and models for the putative alternate state.

**Figure 5 – Figure Supplement 2:**
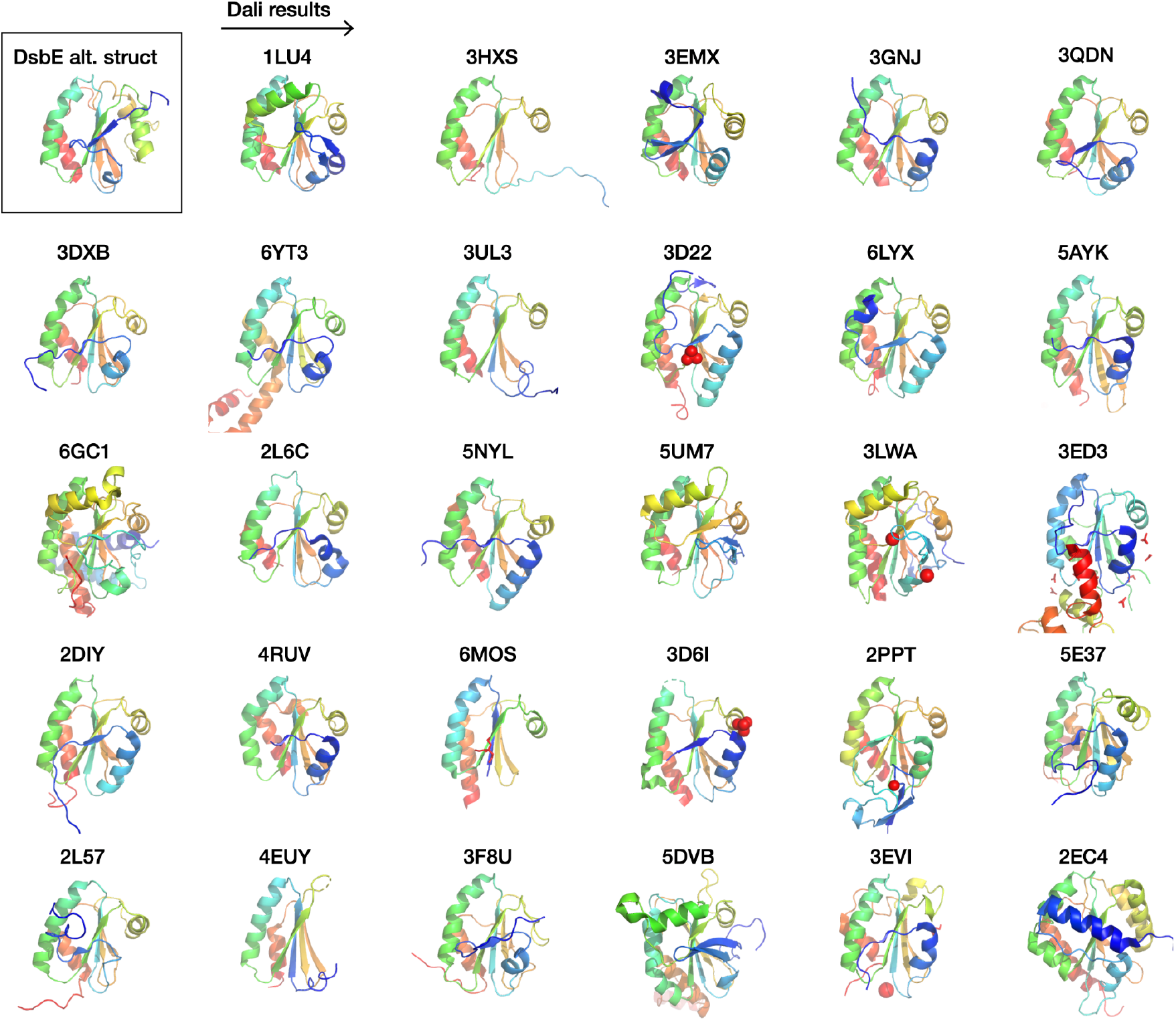
Top candidates for DsbE alternate state from Dali. Screening for the putative alternate state of DsbE in the PDB results in folds that match original thioredoxin fold. Note that the top-ranked hit is the known thioredoxin-like fold of DsbE.

**Figure 6 – Figure supplement 1:**
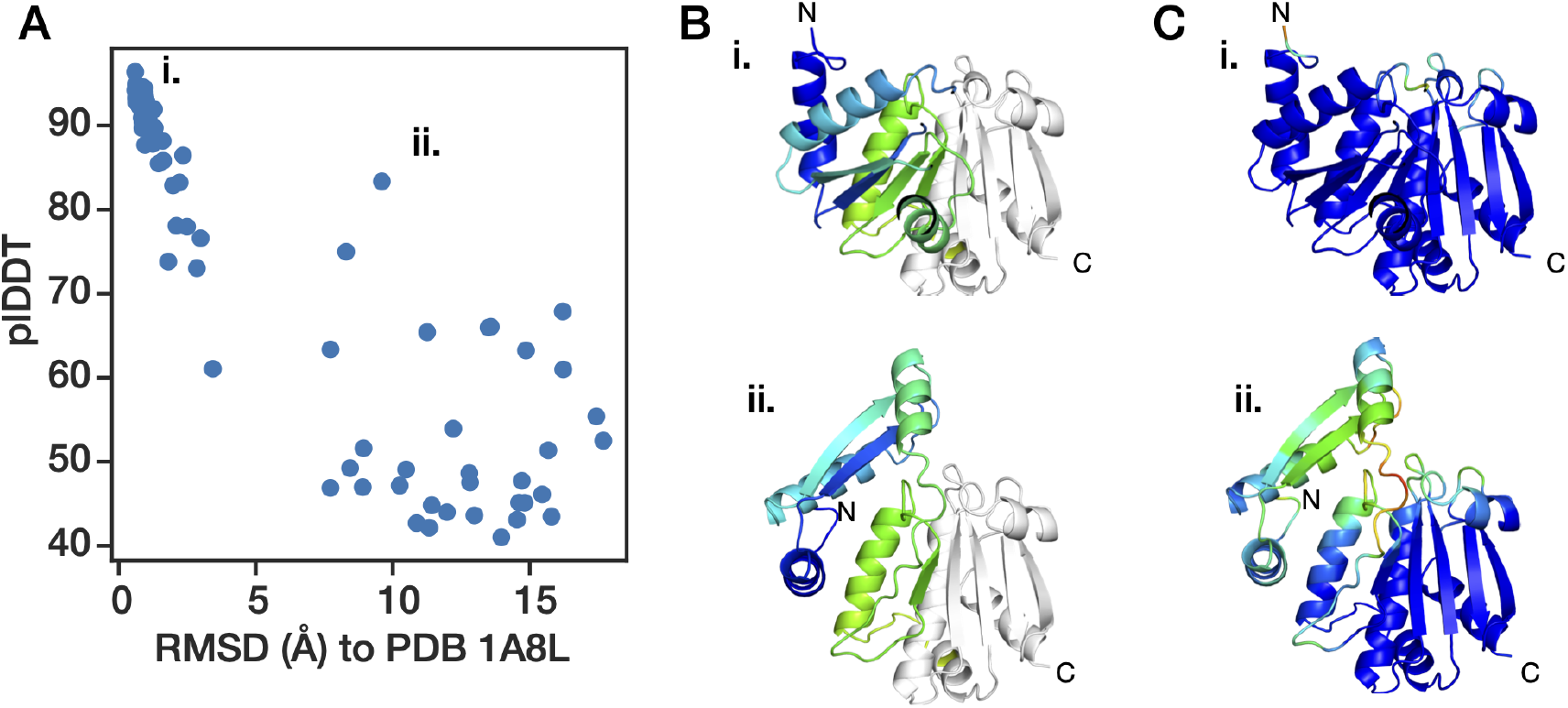
Predicted alternate state for *P. furiosus* oxidoreductase. a) plDDT vs. RMSD to crystal structure 1A8L. b) Models for known state (i) and alternate state (ii), which contains two repacked N-terminal β-strands, colored where the structures diverge. c) Same structures as in (b), colored by plDDT by residue.

